# L-WNK1 is required for BK channel activation in intercalated cells

**DOI:** 10.1101/2021.01.26.428162

**Authors:** Evan C. Ray, Rolando Carrisoza-Gaytan, Mohammad Al-Bataineh, Allison L. Marciszyn, Lubika J. Nkashama, Jingxin Chen, Aaliyah Winfrey, Daniel Flores, Peng Wu, WenHui Wang, Chou-Long Huang, Arohan R. Subramanya, Thomas R. Kleyman, Lisa M. Satlin

**Author notes:** Address all correspondence to: Thomas Kleyman, MD, Renal-Electrolyte Division, University of Pittsburgh, A919 Scaife Hall, 3550 Terrace Street, Pittsburgh, PA 15261. Authors contributed equally to this work (Evan C. Ray, Rolando Carrisoza-Gaytan).

## Abstract

BK channels expressed in intercalated cells (ICs) in the aldosterone-sensitive distal nephron (ASDN) mediate flow-induced K^+^ secretion. In the ASDN of mice and rabbits, IC BK channel expression and activity increase with a high K^+^ diet. In cell culture, the long isoform of the kinase WNK1 (L-WNK1) increases BK channel expression and activity. Apical L-WNK1 expression is selectively enhanced in ICs in the ASDN of rabbits on a high K^+^ diet, suggesting that L-WNK1 contributes to BK channel regulation by dietary K^+^. We examined the role of IC L-WNK1 expression in enhancing BK channel activity in response to a high K^+^ diet. Mice with an IC-selective deletion of L-WNK1 (IC-L-WNK1-KO) and littermate control mice were placed on a high K^+^ (5% K^+^ as KCl) diet for at least 10 days. IC-L-WNK1-KO mice exhibited higher blood K^+^ concentrations ([K^+^]) than controls. BK channel-dependent whole-cell currents in ICs from cortical collecting ducts of high K^+^ fed IC-L-WNK1-KO mice were reduced compared to controls. Six-hour urinary K^+^ excretion in response a saline load was similar in IC-L-WNK1-KO mice and controls. The observations that IC-L-WNK1-KO mice have higher blood [K^+^] and reduced IC BK channel currents are consistent with impaired urinary K^+^ secretion, and suggest that IC L-WNK1 has a role in the renal adaptation to a high K^+^ diet.

---

A moderate daily consumption of K^+^ in humans (∼ 100 mmoles) exceeds the total extracellular K^+^ content of an adult (∼ 70 mmoles) (16). Life-threatening hyperkalemia would certainly occur if not for the initial rapid intracellular buffering of excess K^+^ that enters the extracellular space, and the eventual urinary excretion of K^+^. In the context of a high K^+^ diet, the aldosterone-sensitive distal nephron (ASDN) is a key site of secretion of K^+^ into the tubular lumen, leading to net urinary K^+^ excretion. High flow rates of tubular fluid further enhance K^+^ secretion in the ASDN, increasing urinary K^+^ excretion (11, 22, 26, 30, 33, 52).

An iberiotoxin-sensitive, large conductance, Ca^2+^-and/or stretch-activated K^+^ channel, referred to as BK or K_Ca_1.1, has a critical role in flow-induced K^+^ secretion (FIKS) in the ASDN (6, 7, 33, 46, 52). A high K^+^ environment (in salamanders) or high K^+^ diet (in rabbits and mice) stimulated apical expression of this channel as well as BK channel-mediated K^+^ secretion in the collecting duct (CD) (2, 7, 9, 33, 46, 52). A global knock-out of the mouse BK α or β1 subunit provided support for the role of BK channels in renal K^+^ secretion (39, 40).

ENaC dependent Na^+^ absorption in the ASDN has an important role in K^+^ secretion and FIKS (25, 35), although ENaC independent K^+^ secretion has been observed (14). BK channels are expressed in both principal cells (PCs) and ICs (7, 9, 13, 33, 36). The question of whether BK channels in PCs, ICs or both cell types are responsible for FIKS has been addressed by several groups. There is a growing body of evidence supporting a key role of IC BK channels in FIKS. BKα expression in PCs is concentrated within the cells’ single apical cilium (9). Chemical deciliation of PCs did not impair FIKS in isolated, perfused cortical collecting ducts (CCDs), suggesting that PCs do not have a central role in FIKS. Transition of rabbits to a high K^+^ diet increases BKα subunit expression primarily in ICs, rather than PCs (9). In mice, an IC-selective knockout of the BKα subunit abolished FIKS in isolated, perfused CCDs from these animals and was associated with an increased blood [K^+^], at least in male mice on a high K^+^ diet (7).

Mechanisms by which increases in dietary K^+^ enhance IC BKα expression and BK channel activity remain unclear. Previous studies suggest members of the WNK (with-no-lysine) kinase family are intermediaries in this process. WNK4 inhibited protein and functional expression of the BKα subunit in HEK293 cells (48, 55, 56). In contrast, both the long form of WNK1 (L-WNK1) and a truncated, kidney specific isoform of WNK1 (KS-WNK1) augmented expression of the BKα subunit in cultured cells (HEK293) (28, 50). Furthermore, rabbits fed a high K^+^ diet exhibited enhanced apical expression of L-WNK1 selectively in ICs (50). As L-WNK1 inhibits ROMK (12, 23), a key channel mediating K^+^ secretion in PCs, an IC selective increase in the expression of L-WNK1 has been proposed to have an important role in the renal response to a high K^+^ diet (50). These observations led us to examine the role of L-WNK1 in ICs in the renal adaptive response to a high K^+^ diet. A mouse model with an IC-specific disruption of L-WNK1 (IC-L-WNK1-KO) was generated and characterized. We found that gene disruption of L-WNK-1 in ICs blunted charybdotoxin (ChTX)-sensitive whole cell K^+^ currents in ICs in isolated CCDs from mice on a high K^+^ diet. When placed on a high K^+^ diet, IC-L-WNK1-KO mice had higher blood [K^+^] than littermate controls, suggesting impaired IC-dependent K^+^ excretion.

## Methods

### Animals

Mice were allowed free access to tap water and standard mouse chow (Prolab 5P76 Isopro RMH 3000), which was mixed with water for the IC-L-WNK1-KO mice (see below). All animal breeding, housing, and protocols were approved by the Institutional Animal Care and Use Committees at the University of Pittsburgh, the Icahn School of Medicine at Mount Sinai, and New York Medical College. Mouse facilities were accredited by the American Association for the Accreditation of Laboratory Animal Care. Animals were euthanized in accordance with the National Institutes of Health Guidelines for the Care and Use of Laboratory Animals.

### Generation of IC-specific L-WNK1-KO mice

Mice with loxP sites flanking exon 2 of Wnk1 (L-Wnk1^fl/fl^) in a C57Bl/6 background (53) (Figure 1), and mice expressing the Cre recombinase under control of the promoter for Atp6v1b1, encoding the vacuolar ATPase B1 subunit (B1:Cre mice; C57Bl/6 background) (31), were previously characterized. Deletion of exon 2 leaves KS-WNK1 intact, while ablating L-WNK1 expression in cells expressing the Cre recombinase. To confirm cell-specific Cre expression, mice hemizygous for B1:Cre were crossed with Rosa26^tdTomato^ Cre reporter mice (7). Kidney cortex cryosections were labeled with a polyclonal goat anti-aquaporin-2 (Aqp2) antibody (0.2 μg/ml, sc-9882; Santa Cruz Biotechnology, Santa Cruz CA, USA), and detected using a donkey anti-goat immunoglobulin G conjugated to Cyanine Cy3 (0.5 μg/ml, Jackson ImmunoResearch Laboratories, West Grove, PA, USA). Images were acquired with a Leica TCS SP5 confocal microscope. The Atp6v1b1 promoter selectively drives expression of Cre in Aqp2-negative tubular cells. Occasional staining in Aqp2-positive tubular cells was seen, as previously reported (32) (Figure 2).

**Figure 1.**
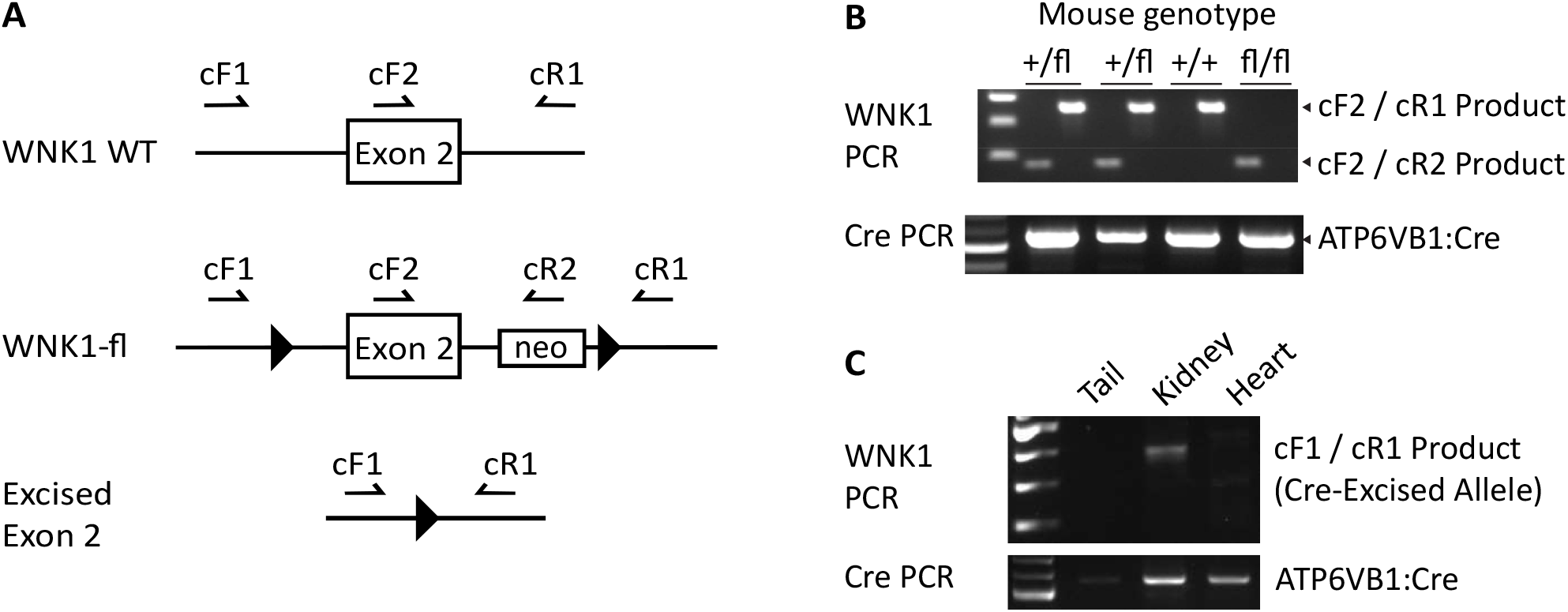
Generation of L-WNK1-IC-KO mice. **A**. Primers used for genotyping and confirmation of L-WNK1 exon 2 excision (not to scale). Sequences are as previously reported (53). **B**. Primers cF2 and cR1 amplify the wild-type (non-floxed) WNK1 locus into a ∼350 base pair PCR product. cF2 and cR2 generate a 180 base-pair product from the floxed allele. **C**. The cF1/cR1 primer pair amplified the excised allele in kidney but not in heart or tail.

**Figure 2.**
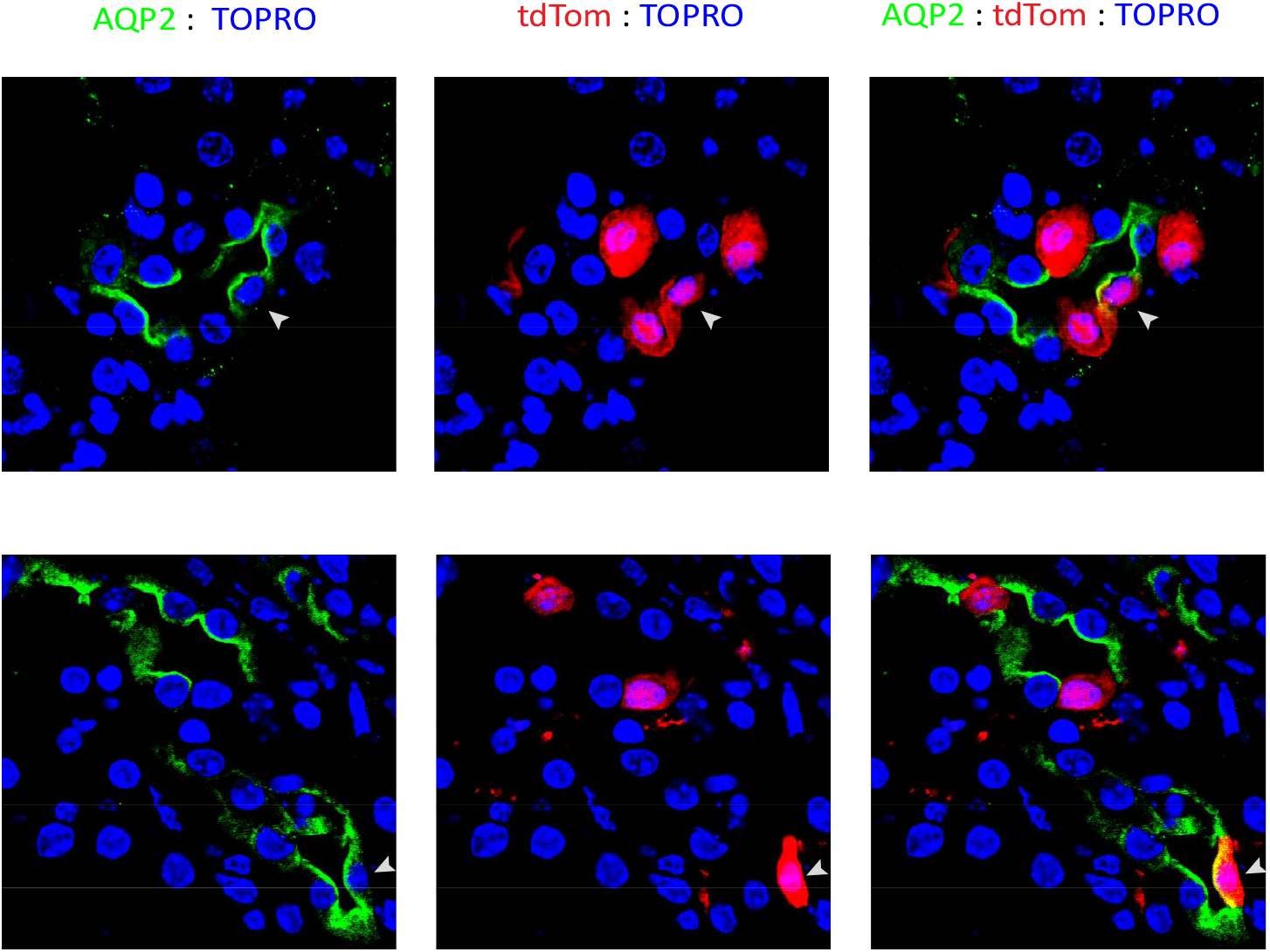
The vacuolar ATPase B1 subunit promoter drives Cre expression predominantly in Aqp2-negative cells. Aqp2 expression is shown in green. tdTomato is shown in red. Nuclei are highlighted with TO-PRO-3 staining. Most Aqp2-negative cells express tdTomato. A minority of Aqp2-expressing cells also express tdTomato (arrowhead). Representative images of kidney sections from 2 mice.

To generate IC-L-WNK1-KO mice, male homozygous floxed L-WNK1 mice were bred with female B1:Cre-expressing mice. Genotyping was performed prior to weaning by PCR amplification, using the primer scheme shown in Figure 1 (primer details in (53)). Female offspring hemizygous for B1:Cre and heterozygous for floxed L-WNK1 were bred with male homozygous floxed L-WNK1 mice to obtain mice homozygous for floxed L-WNK1 and hemizygous for B1:Cre (IC-L-WNK1-KO). Sex-matched littermates that were homozygous for floxed L-WNK1, but without Cre, were used as controls, and mice of both sexes were used for all studies. After weaning, IC-L-WNK1-KO mice failed to thrive on solid food. When these mice were provided chow mixed with water, IC-L-WNK1-KO mice had weights similar to controls, as described in *Results*.

### Blood and Urine Metabolite Measurements

Ten to 13 week-old mice were placed on a control K^+^ gel diet (1% K^+^, TD.88238, Envigo) for a minimum of 2 days, and then switched to a high K^+^ gel diet (5.2% K^+^ as KCl, TD.09075, Envigo) for up to 11 to 13 d. To generate the gel diet, 3 gm agar was dissolved in 250 ml boiling H_2_O. The solution was cooled, 125 gm of powdered control (1% or 5.2% K^+^, TD.88238, Envigo) diet was incorporated, and the slurry was aliquoted into dishes to cool and harden before being placed in cages. After 10 d on the high K^+^ diet, some mice were sacrificed to obtain blood to measure electrolytes. Other mice were placed in a metabolic cage for approximately 1 hr and encouraged to void, and then given 10% vol/wt sterile 0.9% saline via intraperitoneal injection. Mice were replaced in the metabolic cage, and urine was collected over 6 hours. Urine Na^+^ and K^+^ excretion were measured via flame photometry (IL 943, Instrumentation Laboratory, Bedford, MA). The following day, mice were sacrificed under 2-5% isoflurane anesthesia. Thoracotomy was performed, exposing the heart. Blood was aspirated from the cardiac ventricles using a heparinized syringe. An i-STAT handheld analyzer was used to measure electrolytes in whole blood (Abbot Point-of-Care, Princeton, NJ, USA). Plasma aldosterone levels were measured using an aldosterone ELISA kit (Enzo Life Sciences, Farmingdale, NY, USA)

### Immunofluorescence Microscopy

One kidney from each mouse was sliced lengthwise and placed in 4% paraformaldehyde in PBS for 16 h at 4°C. After immersion in 30% sucrose in PBS with 0.02% azide, 5 µm-thick sections were prepared as described previously (4). Immunofluorescence labeling was performed with a rabbit antibody raised against amino acids 2031-2117 of the human L-WNK1 carboxyl-terminus (3 μg/ml, Atlas, HPA059157, Bromma, Sweden) (5), followed by a secondary donkey anti-rabbit Ig antibody coupled to Alexa 488 (3 μg/ml, Jackson ImmunoResearch Laboratories, West Grove, PA). A goat anti-Aqp2 antibody (0.2 μg/ml, as described above) and a secondary donkey anti-goat Ig antibody with conjugated Cy3 (0.5 μg/ml, Jackson ImmunoResearch Laboratories, West Grove, PA) identified PCs. TO-PRO-3 Cy5 was obtained from Molecular Probes (ThermoFisher, Grand Island, NY). Immunolabeled tissues were mounted in mounting medium (Vector Laboratories, Burlingame, CA), and imaged with a confocal laser scanning microscope (Leica TCS SP5, Model upright DM 6000S, Leica Microsystems Inc., Buffalo Grove, IL, USA) using a 63x objective with identical laser settings for all samples.

### Patch-clamp electrophysiology

BK channel activity was assessed via perforated whole-cell recording of ICs in micro-dissected, split open CCDs from IC-L-WNK1-KO and littermate control mice on a high K^+^ diet for 10 days, as previously described (7). The bath solution contained (in mM) 130 K^+^ gluconate, 10 KCl, 0.5 MgCl_2_, 1.5 CaCl_2_, 10 HEPES, pH 7.4, whereas the pipette solution had 130 K^+^ gluconate, 10 KCl, 1 MgCl_2_, 8.72 CaCl_2_, 10 EGTA (1 μM free Ca^2+^), 5 HEPES, pH 7.4. After forming a high-resistance seal (>2 GΩ), the membrane capacitance was monitored until the whole-cell patch configuration was formed. The IC membrane capacitance ranged between 12.5 to 13.5 pF; currents were normalized to a membrane capacitance of 13 pF per cell. Cells were clamped at +60 mV and outward BK currents were identified by adding 100 nM charybdotoxin (ChTx) to the bath solution. Currents, recorded using an Axon 200A patch-clamp amplifier, were low-pass filtered at 1 KHz, and digitized by an Axon interface (Digidata 1440A, Molecular Devices) (27, 55). Data were analyzed using the pCLAMP Software System 9 (Molecular Devices).

### Data analysis

Data were analyzed in Graphpad Prism v. 8.0.2. Removal of outliers was performed using the ROUT method with a ROUT coefficient of 1%. Reported errors and error bars shown all represent standard deviations. Experimental and control data sets were analyzed via two-tailed Student’s t-test. A p value of less than 0.05 was considered statistically significant.

## Results

### IC-specific L-WNK1 knock out (IC-L-WNK1-KO) mice

To clearly define the role of L-WNK1 in enhancing IC-dependent BK expression and facilitating the adaptation to a high K^+^ diet, we generated both mice where L-WNK1 was selectively deleted from ICs (IC-L-WNK1-KO) and littermate controls. Expression of the Cre recombinase in Aqp2-negative tubular epithelial cells was confirmed by crossing B1:Cre mice with Ai9 tdTomato-expressing Cre-reporter mice (29). Tubules from these mice demonstrated robust expression of tdTomato in Aqp2-negative cells within Aqp2-expressing tubules (Figure 2). A minor fraction of Aqp2-expressing cells also expressed tdTomato, consistent with previous work suggesting that B1 promoter-driven Cre expression was occasionally seen in PCs (31).

Genotyping was performed as previously described (53) and recombination between L-WNK1 loxP sites flanking exon 2 was confirmed by PCR (Figure 1). A recombinant band was observed in kidney, but not tail or heart of IC-L-WNK1-KO mice. IC-L-WNK1-KO mice were born at a frequency of approximately 20% lower than expected, gained weight when chow was mixed with water (see *Methods*), developed normally, and had no gross morphologic abnormalities. We examined WNK1 expression in kidney sections from IC-L-WNK1-KO and littermate control mice maintained on a high K^+^ diet for at least 10 days, using an antibody directed against the C-terminus of WNK1 (which recognizes both L-WNK1 and KS-WNK1). We did not observe differences in WNK1 expression in Aqp2-negative cells within Aqp2-expressing tubules obtained from IC-L-WNK1-KO and littermate control mice (not shown). The lack of difference in WNK1 expression in ICs from IC-L-WNK1-KO and littermate control mice may reflect expression of KS-WNK1 in ICs.

### IC-L-WNK1-KO mice exhibited an elevated blood [K^+^] on a high K^+^ diet

IC-L-WNK1-KO mice developed normally and resembled control mice in terms of body weight (Figure 3 and Table 1). At 10-13 weeks of age mice transitioned to a high K^+^ diet. Whole blood electrolytes were assessed after 10-13 days on the high K^+^ diet. Blood K^+^ was significantly higher in the IC-L-WNK1-KO mice (5.4 ± 1.0 mM, n=26), compared to littermate controls (4.8 ± 0.6 mM, n=19, p<0.05, Figure 3 and Table 1). When analyzed by sex, significant differences in blood [K^+^] were not observed (not shown). Similar levels of blood Na^+^, Cl^-^, total CO_2_, Ca^2+^, urea nitrogen, and hemoglobin were observed in IC-L-WNK1-KO mice and littermate controls. Furthermore, plasma aldosterone levels did not differ between IC-L-WNK1-KO mice and controls (Table 1).

**Table 1.**

|  | **L-WNK1<sup>fl/fl</sup>**<br>**Control** | **L-WNK1<sup>fl/fl</sup>**<br>**B1:Cre** | **p** |
| --- | --- | --- | --- |
| **Blood** |  |  |  |
| K<sup>+</sup> (mmol/L) | 4.8 ± 0.6 (19) | 5.4 ± 1.0 (26) | 0.037 |
| Na<sup>+</sup> (mmol/L) | 146 ± 2 (19) | 146 ± 2 (26) | NS |
| Cl<sup>-</sup> (mmol/L) | 115 ± 4 (19) | 116 ± 4 (26) | NS |
| tCO<sub>2</sub> (mmol/L) | 21.2 ± 2 (19) | 20.3 ± 3 (25) | NS |
| Ionized Ca<sup>2+</sup> (mmol/L) | 1.26 ± 0.12 (19) | 1.26 ± 0.11 (26) | NS |
| Urea Nitrogen<br>(mg/dL) | 20 ± 4 (19) | 22 ± 4 (26) | NS |
| Hemoglobin (mg/dL) | 12.7 ± 1.0 (19) | 12.4 ± 1.1 (26) | NS |
| **Plasma** |  |  |  |
| Aldosterone | 1969 ± 2165 (16) | 1779 ± 1978 (16) | NS |
| **Urine** |  |  |  |
| UNaV̇ (μmol/hr) | 29.6 ± 8.2 (19) | 32.2 ± 9.7 (15) | NS |
| UKV̇ (μmol/hr) | 26.5 ± 11.6 (19) | 28.1 ± 9.2 (15) | NS |
| **Mouse Weights** |  |  |  |
| Females (gm) | 24.3 ± 3.4 (9) | 23.4 ± 1.8 (7) | NS |
| Males (gm) | 29.3 ± 4.0 (11) | 29.3 ± 3.5 (15) | NS |

Blood and urine measurements were obtained in IC-L-WNK1-KO and littermate control mice after 10 days on a high K<sup>+</sup> diet, as described under *Methods*. Differences were assessed using a two-tailed Student's t-test. NS=not significant.

**Figure 3.**
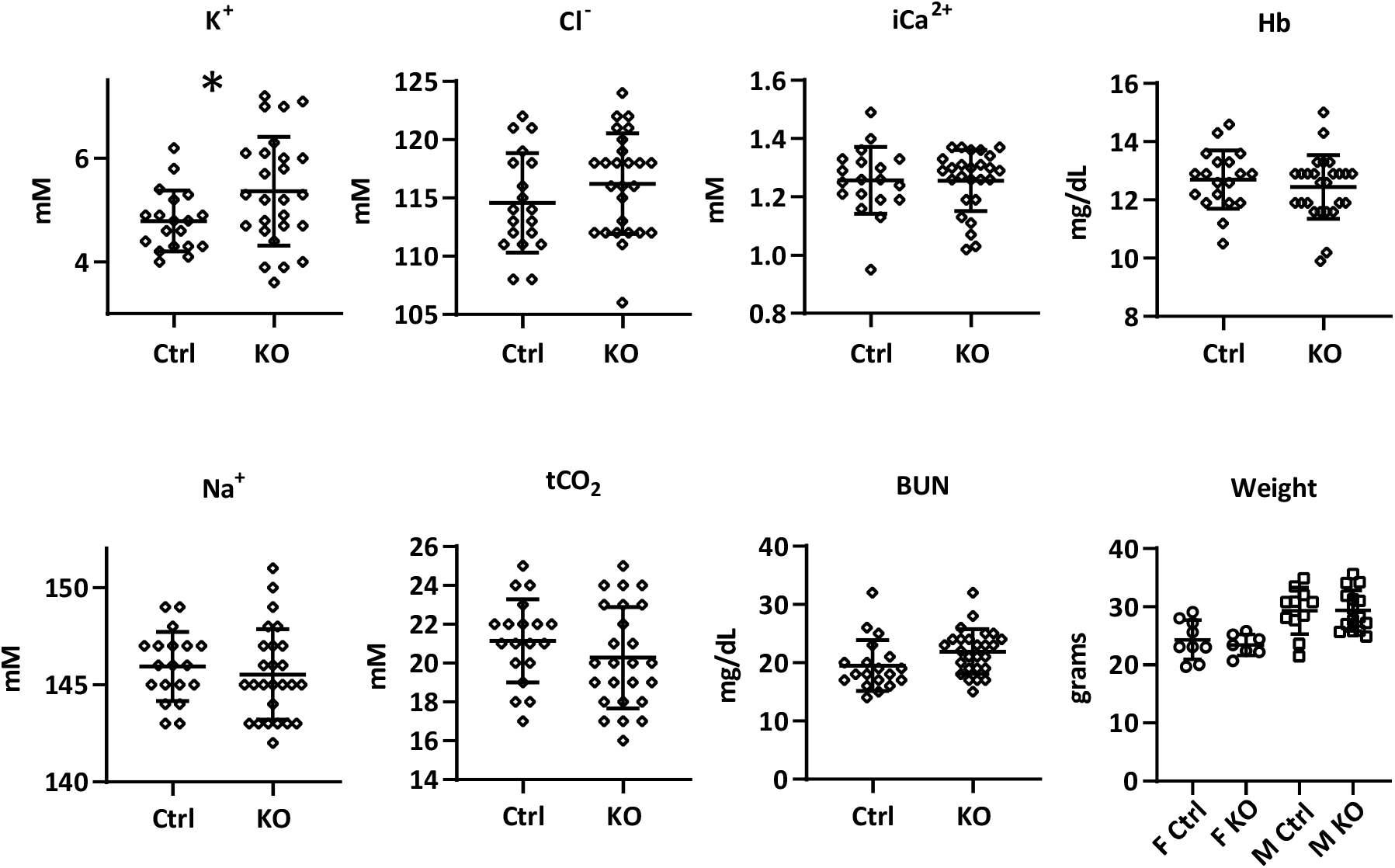
L-WNK1-IC-KO mice on a high K^+^ diet exhibited higher blood [K^+^] than littermate controls. Other electrolytes, hemoglobin (Hb), and blood urea nitrogen (BUN) did not differ. Weights were similar between L-WNK1-IC-KO mice and controls. Means ± SD are shown. * p<0.05, IC-L-WNK1-KO vs. littermate controls, two-tailed Student’s t-test.

### IC BK channel currents are reduced in IC-L-WNK1-KO mice

We next examined whether BK channel activity in ICs was reduced in IC-L-WNK1-KO mice, when compared to littermate controls. Perforated whole cell patch clamp recordings of K^+^ currents were performed on ICs clamped at +60 mV in isolated tubules from IC-L-WNK1-KO and littermate control mice on a high K^+^ diet for 10 or more days. Currents were normalized to a membrane capacitance of 13 pF per cell. Charybdotoxin-sensitive currents, reflecting BK channel-mediated currents (8), were significantly reduced in ICs from IC-L-WNK1-KO mice (452 ± 73 pA, n=8) compared to littermate controls (691 ± 91 pA, n=8, p<0.001, two-tailed Student’s t-test, Figure 4).

**Figure 4.**
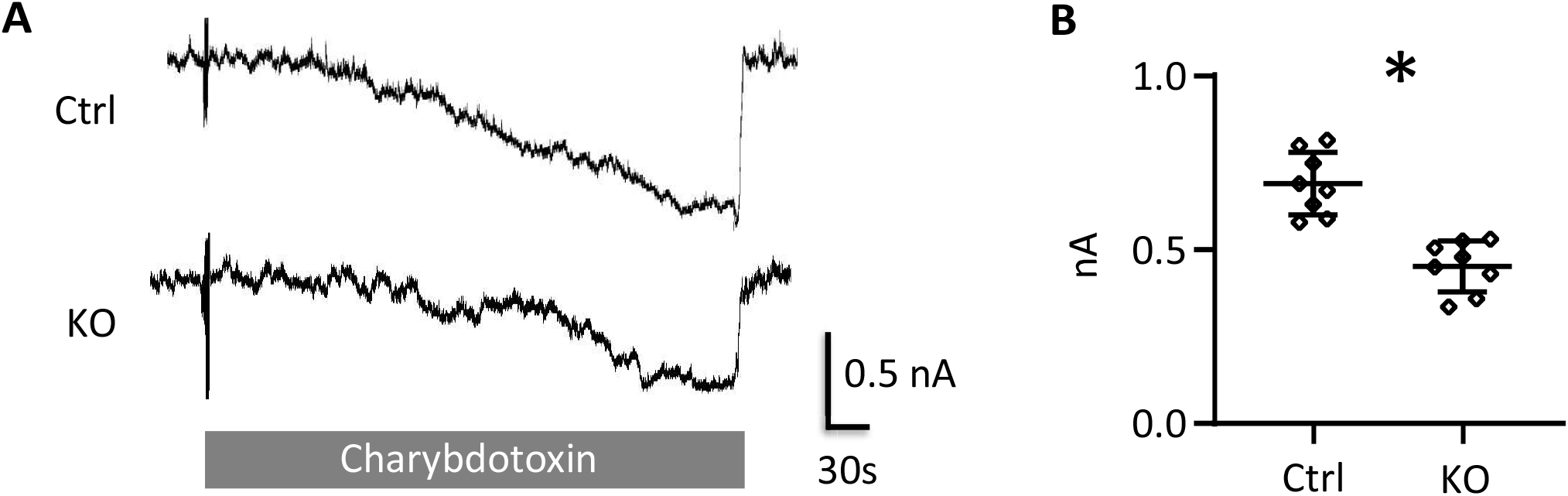
Charybdotoxin-sensitive currents (indicative of BK channel activity) were reduced in intercalated cells (ICs) from L-WNK1-IC-KO mice on a high K^+^ diet. Perforated whole-cell current tracings were collected from ICs in micro-dissected tubules from L-WNK1-IC-KO or littermate control mice on a high K^+^ diet. Means ± SD are shown. * p<0.05, IC-L-WNK1-KO vs. littermate controls, two-tailed Student’s t-test.

### Urinary Na^+^ and K^+^ excretion

As BK channels in ICs mediate FIKS, we examined whether high K^+^ fed IC-L-WNK1-KO mice exhibited diminished urinary K^+^ excretion 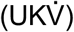 in the context of an intraperitoneal-administered bolus of normal saline (10% of body weight), when compared to littermate controls. Urine was collected over a 6 h period, and 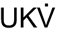 and 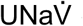 were determined. IC-L-WNK1-KO and littermate control mice exhibited similar 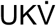 (controls: 26.5 ± 11.6 µmol/hr, n = 15; IC-L-WNK1-KO: 28.1 ± 9.2 µmol/hr, n = 19, p = not significant) and similar 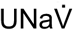 (controls: 29.6 ± 8.2 µmol/hr, n = 15; IC-L-WNK1-KO: 32.2 ± 9.7 µmol/hr, n = 19, p = not significant; Figure 5 and Table 1).

**Figure 5.**
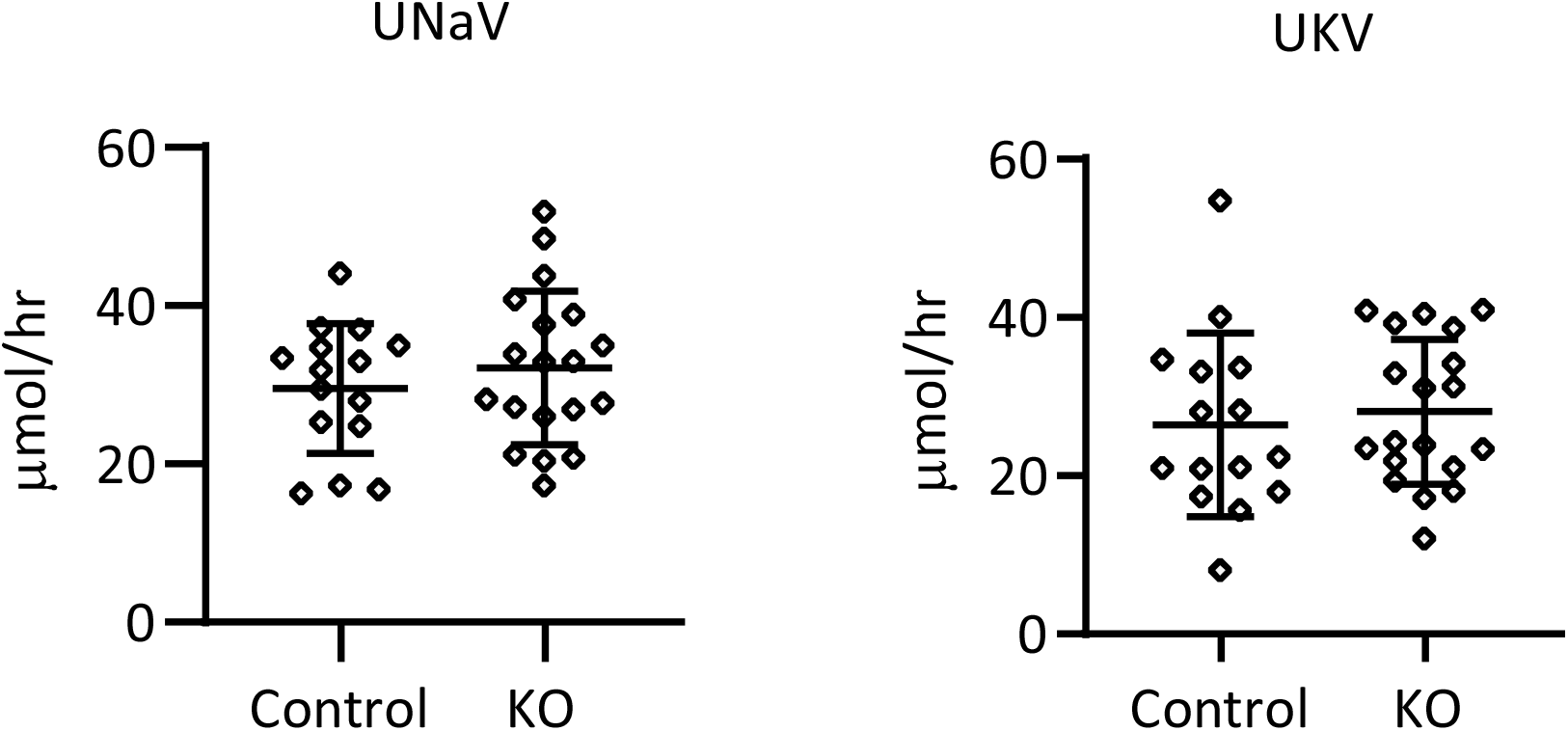
Urinary K^+^ and Na^+^ excretion in response to a fluid bolus did not differ between L-WNK1-IC-KO and littermate control mice. Mice were placed on a high K^+^ diet for at least 10 days and given a 10% body weight injection of 0.9% saline. Urine was collected in metabolic cages over 6 hours and used to calculate hourly excretion. Means ± SD are shown. p=not significant, IC-L-WNK1-KO vs. littermate controls, two-tailed Student’s t-test.

## Discussion

The WNK family of protein kinases has a key role in coordinating the regulated absorption of Na^+^ and secretion of K^+^ in the ASDN (17, 20). In the distal convoluted tubule (DCT), both L-WNK1 and WNK4 activate a signaling cascade involving Sterile20-related proline-alanine-rich kinase and oxidative stress responsive kinase-1 (SPAK and OSR1) to eventually activate the thiazide-sensitive Na-Cl cotransporter (NCC), enhancing Na^+^ and Cl^-^ absorption (17, 20). In more distal aspects of the nephron, WNK4 has been shown to inhibit ENaC (41, 42, 54), whereas KS-WNK1 activates ENaC (18, 34). Interestingly, both WNK4 and L-WNK1 activate SGK1 (19), a well characterized activator of ENaC (3). Furthermore, SPAK has been shown to activate ENaC (1). Most of this work has been performed in heterologous expression systems. Studies of mouse models where specific WNKs have been genetically deleted in PCs are needed to clarify how specific WNKs modulate ENaC *in vivo*.

WNKs have differing roles in regulating K^+^ secretory channels. ROMK is inhibited by L-WNK1 and WNK4 (12, 21, 23, 24, 42). In heterologous expression systems, WNK4 inhibits BK channels (48, 55, 56). In contrast, both L-WNK1 and KS-WNK1 increase both surface and functional expression of BK channels, and its kinase activity is not required for BK channel activation (28, 50). Previous work has suggested that L-WNK1 activates BK channels by inhibiting ERK1/2-dependent channel degradation (28).

BK channels are responsible for FIKS (6, 7, 33, 46, 52). While BK channels are expressed in both PCs and ICs, our recent work suggests that BK channels in ICs mediate FIKS (7). Our current work suggests L-WNK1 has a role in activating BK channels in ICs in mice fed a high KCl diet. Specifically, whole IC BK currents, as measured by the charybdotoxin-sensitive component of the K^+^ current, were significantly reduced in IC-L-WNK1-KO mice fed a high KCl diet, compared to littermate controls. Consistent with reduced BK channel activity, IC-L-WNK1-KO mice on a high KCl diet had significantly higher blood [K^+^] compared to littermate controls. We did not observe a sex difference regarding blood [K^+^], as we observed with IC-BK α subunit knockout mice where males selectively exhibited an increased blood [K^+^] on a high K^+^ diet (7). Interestingly, the magnitude of kaliuresis observed in response to volume expansion was similar in IC-L-WNK1-KO and control mice, suggesting compensatory mechanisms in the IC-L-WNK1-KO mice were present to maintain kaliuresis. These mechanisms remain to be determined.

We previously found that L-WNK1 expression is selectively enhanced in ICs in response to a high K^+^ diet (50). Why this occurs is still unclear. However, our data suggest that an IC specific increase of L-WNK1 expression should enhance BK channel activity, as BK channel activity was reduced in ICs from IC-L-WNK1-KO mice (Fig 5). There are a number of cellular factors that could influence L-WNK1 expression and activity in response to an increase in dietary K^+^. For example, L-WNK variants express PY motifs that are binding sites for the ubiquitin ligase Nedd4-2, targeting L-WNK1 for ubiquitination and degradation (43). Aldosterone-dependent activation of the SGK1 in ICs could prevent Nedd4-2-dependent L-WNK1 degradation. WNKs are inhibited by increases in intracellular [Cl^-^] (10, 38), although it is not known whether a high K^+^ diet is associated with a change in intracellular [Cl^-^] in ICs that could affect L-WNK1 activity. It is interesting that aldosterone binding and subsequent signaling in ICs is dependent on dephosphorylation of Ser483 on the mineralocorticoid receptor, and hyperkalemia enhances Ser483 phosphorylation via unc-51-like kinase 1 (ULK1) (44, 45). Previous work suggests that WNK1 reduces expression of ULK1 (15), raising the possibility of a positive feedback loop. IC MR knockout mice exhibit hyperkalemia (37), proposed to be due to reduced ENaC activity, in turn reducing the driving force for K^+^ secretion. It could also reflect dampened BK channel-mediated K^+^ excretion, as administration of the MR antagonist spironolactone to rats reduced BK α subunit abundance (51).

BK channel mediated currents were moderately reduced in ICs from IC-L-WNK1-KO mice, as compared to the virtual absence of BK channel mediated currents in ICs from mice with a genetic deletion of BK α subunits in ICs (7). These observations are consistent with studies that have identified cellular mechanisms and pathways, in addition to L-WNK1, by which a high K^+^ diet activates BK channel expression and activity. For example, a high K^+^ diet stimulates the expression of CYP2C23 and increases the 11,12-epoxyeicosatrienoic acid concentrations in the CCD, which activates BK channels and FIKS (47). A high K^+^ diet increases expression and activity of HO-1, and CO generated by HO-1 activates BK channels in the CCD by nitric oxide synthase- and protein kinase G-independent mechanisms (49).

In summary, our results demonstrate that L-WNK1 has a role in BK channel activation in ICs in response to a high K^+^ diet. Furthermore, the increased blood [K^+^] in IC-L-WNK1-KO suggests that this signaling pathway has an important role in the adaptation to a high K^+^ diet.

## Acknowledgments

This work was supported by grants from the National institutes of Health, including DK038470 (TRK and LMS), DK110332 (ECR), DK109038 (MAB), DK054983 (WHW), DK111542 (CLH), DK098145 and DK119252 (ARS), and P30 DK079307 (TRK, ARS and LMS). We thank Drs. Donald Kohan and Raoul Nelson at the University of Utah for the B1:Cre mice.

